# Effect of nandrolone decanoate on healing of experimentally induced radial nonunion in rabbits (*Oryctolagus cuniculus*)

**DOI:** 10.1101/2021.03.15.435445

**Authors:** Adrielly Dissenha, Bruno Watanabe Minto, Karina Calciolari, Laís Fernanda Sargi, Lismara Castro do Nascimento, Fabiana Del Lama Rocha, Julián Andrés Sanjuán Galindez, Dayvid Vianêis Farias de Lucena, Luis Gustavo Gosuen Gonçalves Dias

## Abstract

The aim of this study was to evaluate the effect of nandrolone decanoate (ND) on treatment of bone nonunion in the radius of rabbits. Thirty-one, young adult, New Zealand White rabbits (Oryctolagus cuniculus) were allocated to one of four groups: nandrolone males (NMG), nandrolone females (NFG), placebo males (NPM), and placebo females (NPF). After bone nonunion of a 10 mm ostectomy of the radius was confirmed (45 days after surgery), the animals in the NMG and NFG groups received 10 mg/kg ND intramuscular once a week for four weeks, while placebo groups received intramuscular 0.9% NaCl solution. Radiographic, histopathologic, and densitometric parameters (DXA) were used to compared groups.

**Results:** No significant differences were observed radiographically. However, ND groups showed greater area (P=0.0258) and BMC (P=0.0140) in the densitometric evaluation. Histologically, the placebo group showed a predominance of primary bone tissue. Whereas, lamellary organizations of secondary bone and the presence of fibrocartilage were found in the ND group (P =0.006). In conclusion, ND promoted bone regeneration after the creation of a large defect in the radius of rabbits.

## INTRODUCTION

Delayed union and nonunion are common complications of osteosynthesis of long bones in small animal practice. There are many factors resulting in nonunion, including mechanical failures (excessive movement and/or the presence of large bone defects), and damage to the vascularization of the fracture site (1,2). A significant challenge in the treatment of nonunion is the bioactivation of the fracture environment and the filling of critical defects. Classically, the use of bone grafts, bone marrow, and other activation factors has been used; despite treatment a significant proportion of nonunions fail to heal, especially in small dogs (3,4).

Steroids are known to affect bone health and there has been increasing interest in the effect of hormones on bone regeneration. *In vitro*, osteoblastic cells may be stimulated by androgens. Several studies have looked at the use of anabolic steroids as adjuncts to bone repair. Although their mechanisms of action have not been elucidated, they are believed to stimulate osteoblastogenesis, inhibit the action of osteoclasts, and accelerate bone healing. Nandrolone decanoate (ND) is an anabolic steroid used, among other indications, for prevention and treatment of osteopenia as well as to increase bone mass. However, its use *in vivo* requires further investigation (5,6).

In dogs, steroids have been used in the management of osteotomies for tibial tuberosity advancement, and the use of ND and autologous bone grafts, separately and in combination, have been compared; the results demonstrated that the concomitant use of ND and autologous bone grafts decreased the time to bone healing (6). Another study evaluated the use of ND in bone healing of tibial fractures induced in rabbits, and better osteoblastic and collagen proliferation were observed (7).

The aim of this study was to conduct radiographic, histopathologic, and densitometric evaluations on the use of ND in the treatment of, experimentally induced, unviable bone nonunion in rabbits. We hypothesized that the use of ND would correlate with faster and more effective bone regeneration.

## MATERIALS AND METHODS

### Ethics Committee

This study was carried out following the international animal welfare standards after approval by the Ethical Committee of the State University of São Paulo (UNESP) - Jaboticabal / SP (protocol no. 019155).

Thirty-one, young adult (160 to 170 days old), New Zealand White rabbits (*Oryctolagus cuniculus*) were used. They were entire and weighed 3.8 kg (± 0.3). Fifteen were males and 16 were females. Males and females were separately randomly allocated to either treatment or placebo groups, creating four groups: nandrolone males (NMG), nandrolone females (NFG), placebo males (NPM), and placebo females (NPF). One animal from NPM group died of unknown causes on the 75th day.

### Experimental groups

The animals in the NMG and NFG groups received 10 mg/kg ND intramuscularly once a week for four weeks. The PMG and PFG groups received intramuscular 0.9% NaCl solution once a week for 4 weeks. Treatments were started after radiographic confirmation of bone nonunion (45 days after the ostectomy). Radiographic, histopathologic, and densitometric parameters (DXA) were evaluated.

### Surgical procedure

All rabbits underwent the same anesthetic protocol, were positioned in the right lateral decubitus position with the leg extended laterally, and prepared for sterile surgery. A cutaneous incision of approximately 2.5 cm was made in the craniomedial region of the radius. A 10 mm diaphyseal ostectomy of the radius was performed using an oscillatory saw. Local irrigation with 0.9% NaCl solution was used to prevent overheating of the bone. The bone segment was removed with its periosteum, creating a 10 mm bone gap. Soft tissues were routinely sutured with Sultan 4-0 nylon. All surgical procedures were performed by a single surgeon, with experience in general surgery and veterinary orthopedics.

The animals received enrofloxacin (10 mg/kg every 24 hours for 7 days subcutaneously), tramadol hydrochloride (5 mg/kg every 12 hours for 3 days subcutaneously) and meloxicam (1 mg/kg single dose subcutaneously) postoperatively.

### Radiographic evaluation

All rabbits underwent radiographic examination immediately after surgery and on 15, 30, and 45 days postoperatively, until the nonunion was confirmed. ND and placebo treatments began when nonunion was confirmed. The animals were than radiographically evaluated at 60, 75, and 90 days postoperatively (15, 30 and 45 days after confirmation of the nonunion). A double-blind evaluation of the radiographic images was performed by two experienced evaluators.

Bone nonunion was scored before and after ND / placebo administration. Periosteal reaction (PR), volume of bone callus (VBC), and the quality of the bone bridge (QBB) between the fragments were the semi-quantitative parameters used for the evaluation - adapted from previous studies (8,9). The means of the recorded values were used for analysis.

### Bone densitometry and histopathologic evaluation

The animals were euthanized on the 90^th^ day postoperatively using midazolam (1 mg/kg, IM) and propofol (dose-dependent) followed by an intravascular injection of 4 mL of potassium chloride to induce cardiac arrest. Immediately after euthanasia, bone densitometry examination of the dissected radius and ulna was performed. The bones were weighed and measured with a standard millimeter tape. The equipment was calibrated with the use of a phantom, with specific measures for bone analysis, and the bone was then placed on the table and an area of 3 cm was scanned. Bone mineral composition content (g) and bone mineral density (g/cm2) measurements were obtained.

Afterwards, a segment of the radius was collected 10 mm proximally and 10 mm distally from the bone failure interfaces. Bone fragments were fixed in a 10% formalin solution buffered with sodium acid phosphate monohydrate and anhydrous disodium phosphate (pH 7.4), decalcified, embedded in paraffin and stained by the Hematoxylin and Eosin (HE) technique.

The histologic slides were evaluated for three factors: quality of bone neoformation, quantity of bone neoformation and type of bone tissue formed. Quality of bone neoformation, was scored on a scale 0 = no periosteal reaction, 1 = moderate periosteal response, 2 = intense periosteal response, 3 = disorganized neoformed bone tissue, 4 = neoformed bone tissue with moderate organization, and 5 = Neoformed bone tissue with advanced organization. Bone neoformation was scored on the scale where 0 = no bone neoformation or it was not possible to assess, 1 = mild bone neoformation, 2 = mild / moderate bone neoformation, 3 = moderate bone neoformation, 4 = moderate/intense bone neoformation, and 5 = intense new bone formation. The type of bone tissue was evaluated according to the scale where 1 = prevalence of primary bone tissue and 2 = prevalence of secondary bone tissue. (Table 1). All the evaluations were performed by an experienced pathologist.

**Table 1.**
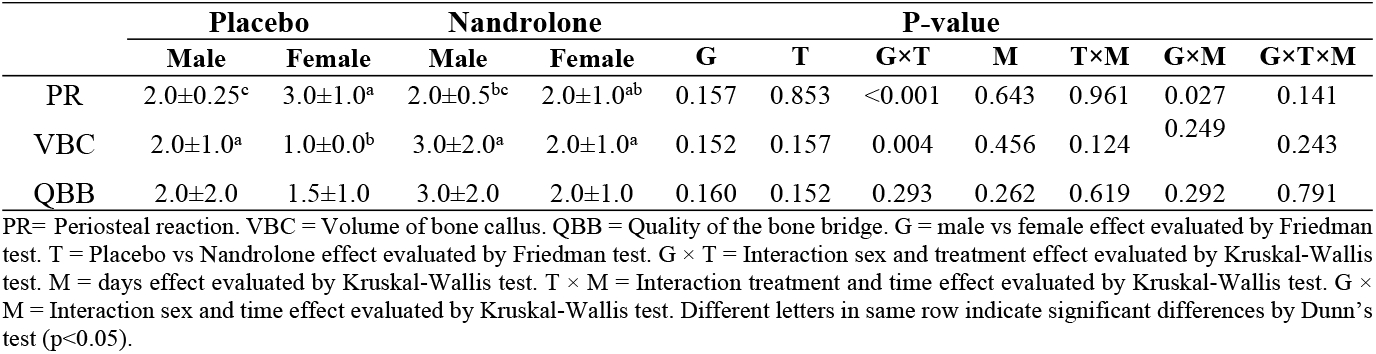
Median ± IQR of radiographic evaluation in rabbits treated with ND versus placebo.

|  | Placebo |  | Nandrolone |  | P-value |  |  |  |  |  |  |
| --- | --- | --- | --- | --- | --- | --- | --- | --- | --- | --- | --- |
|  | Male | Female | Male | Female | G | T | G×T | M | T×M | G×M | G×T×M |
| PR | 2.0±0.25<sup>c</sup> | 3.0±1.0<sup>a</sup> | 2.0±0.5<sup>bc</sup> | 2.0±1.0<sup>ab</sup> | 0.157 | 0.853 | <0.001 | 0.643 | 0.961 | 0.027 | 0.141 |
| VBC | 2.0±1.0<sup>a</sup> | 1.0±0.0<sup>b</sup> | 3.0±2.0<sup>a</sup> | 2.0±1.0<sup>a</sup> | 0.152 | 0.157 | 0.004 | 0.456 | 0.124 | 0.249 | 0.243 |
| QBB | 2.0±2.0 | 1.5±1.0 | 3.0±2.0 | 2.0±1.0 | 0.160 | 0.152 | 0.293 | 0.262 | 0.619 | 0.292 | 0.791 |

PR= Periosteal reaction. VBC = Volume of bone callus. QBB = Quality of the bone bridge. G = male vs female effect evaluated by Friedman test. T = Placebo vs Nandrolone effect evaluated by Friedman test. $G \times T$ = Interaction sex and treatment effect evaluated by Kruskal-Wallis test. M = days effect evaluated by Kruskal-Wallis test. $T \times M$ = Interaction treatment and time effect evaluated by Kruskal-Wallis test. $G \times M$ = Interaction sex and time effect evaluated by Kruskal-Wallis test. Different letters in same row indicate significant differences by Dunn's test ( $p < 0.05$ ).

### Statistical analysis

Statistical analyzes were performed using R Software version 3.6.3 (R Core Team, 2015). Statistical significance for all tests was considered when P ≤ 0.05.

Bone densitometry was evaluated by analyzes of variance (ANOVA) as an A × B Factorial in a randomized design, considering as fixed effects the factor A corresponding to the sex (Male or Female), factor B treatment (Placebo or Nandrolone), factor interactions (sex× treatment), group errors, random animal effects, and residues corresponding to the model. This ANOVA was performed after verification of mathematical assumptions (Shapiro-Wilk test and Bartlett test), and Tukey’s post-hoc was applied when ANOVA indicated a significant difference between means.

Data from both the radiographic and the histologic evaluations were compared among male and female groups, and placebo and Nandrolone groups by a Friedman nonparametric test, and the interaction among groups and evaluation days were evaluated by a Kruskal-Wallis rank test, and Dunn’s Multiple Comparison post-hoc was applied to the significant difference between medians. The presence of cartilage and medullary spaces observed in the histologic evaluation was compared between groups using Fisher’s exact test.

## RESULTS

The females in the placebo group had greater Periosteal Reaction than the males in the Placebo group and the Nandrolone group (Table 1; P <0.001).

Females in both groups at 60 and 90 days had greater Periosteal Reaction than males in both groups at days 45 and 60 (Fig. 1; P = 0.027). The Volume of bone callus and Quality of the bone bridge were not influenced by the groups, evaluation time or their interactions (P> 0.05; Table 1).

**Fig. 1).**
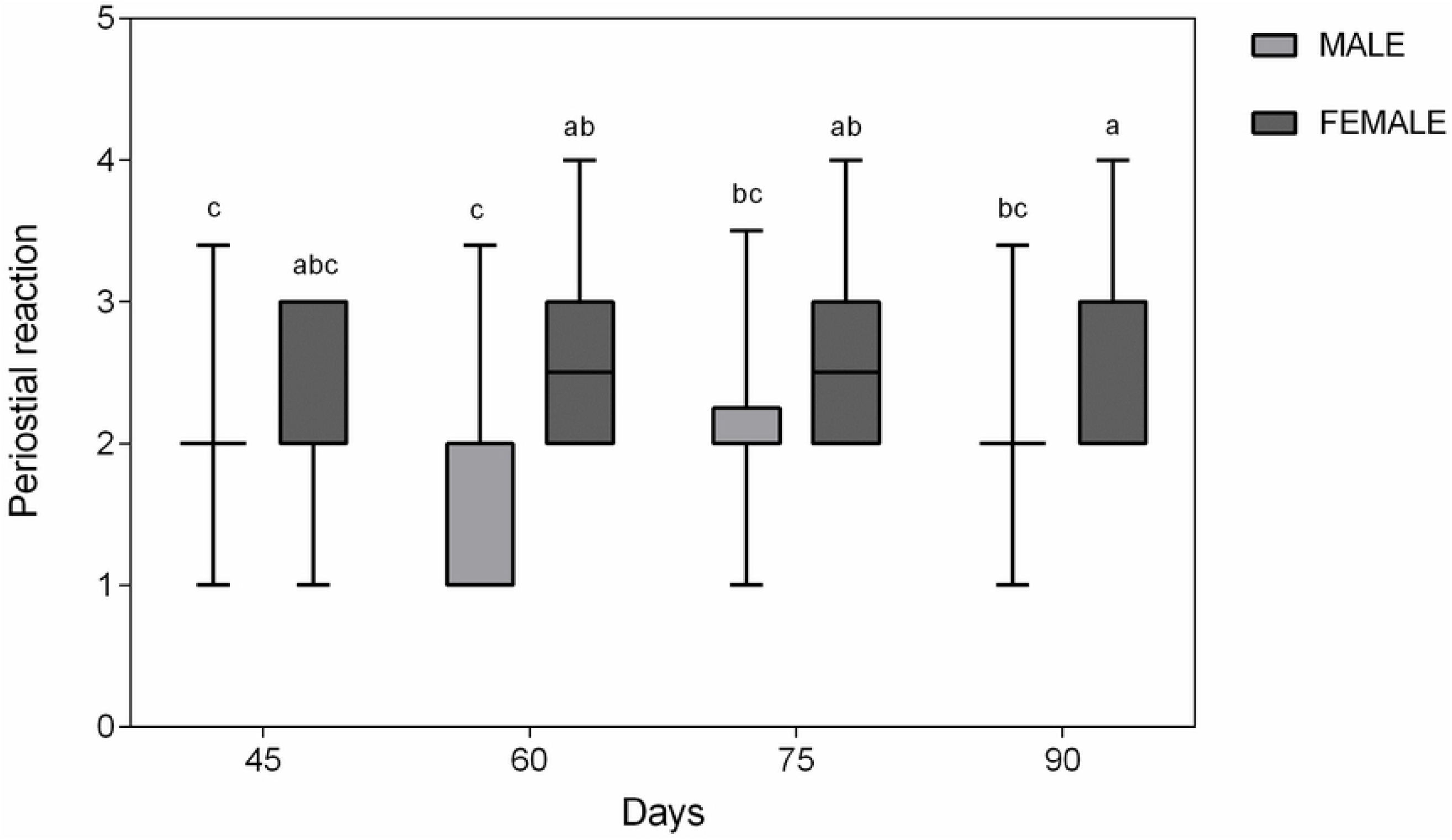
Median and interquartile range of periosteal reaction evaluation by sex and days of evaluation. Different letters indicate significant differences by Dunn’s test (P = 0.027).

The area (cm^2^) and BMC were similar between females and males (P> 0.05; Table 2). However, the groups receiving Nandrolone had a larger area (P = 0.0258; Fig. 2) and a greater BMC (P = 0.0140; Fig. 3) than the Placebo groups. There was an effect of the interaction between sex and treatment in the BMD (g / cm2) (Table 2; P = 0.0192); lower BMD was found in females in the placebo group when compared to the other groups.

**Table 2.** Mean ± SD of Bone densitometry evaluation in rabbits treated with ND versus placebo.

|  | Placebo |  | Nandrolone |  | P-value |  |  |
| --- | --- | --- | --- | --- | --- | --- | --- |
| | Male | Female | Male | Female | G | T | G × T |
| Area, $\text{cm}^2$ | 3.36±0.40 | 3.68±0.45 | 3.81±0.29 | 3.88±0.39 | 0.161 | 0.025 | 0.383 |
| BMC, g | 0.64±0.09 | 0.61±0.09 | 0.71±0.13 | 0.77±0.14 | 0.635 | 0.014 | 0.339 |
| BMD, $\text{g/cm}^2$ | 0.19±0.01<sup>a</sup> | 0.16±0.01<sup>b</sup> | 0.18± 0.02<sup>a</sup> | 0.20± 0.02<sup>a</sup> | 0.533 | 0.091 | 0.019 |
| Area L, $\text{cm}^2$ | 0.78±0.39 | 0.71±0.36 | 0.89±0.18 | 0.74±0.44 | 0.408 | 0.588 | 0.762 |
| BMC L, g | 0.13±0.02 | 0.09±0.01 | 0.15±0.02 | 0.08±0.01 | 0.031 | 0.862 | 0.322 |
| BMD L, $\text{g/cm}^2$ | 0.16±0.03 | 0.13±0.03 | 0.17±0.04 | 0.13±0.06 | 0.046 | 0.738 | 0.995 |

G = male vs female effect. T = Placebo vs Nandrolone effect. G × T = Interaction sex and treatment effect. Different letters in same row indicate significant differences by Tukey test ( $p < 0.05$ ).

**Fig. 2).**
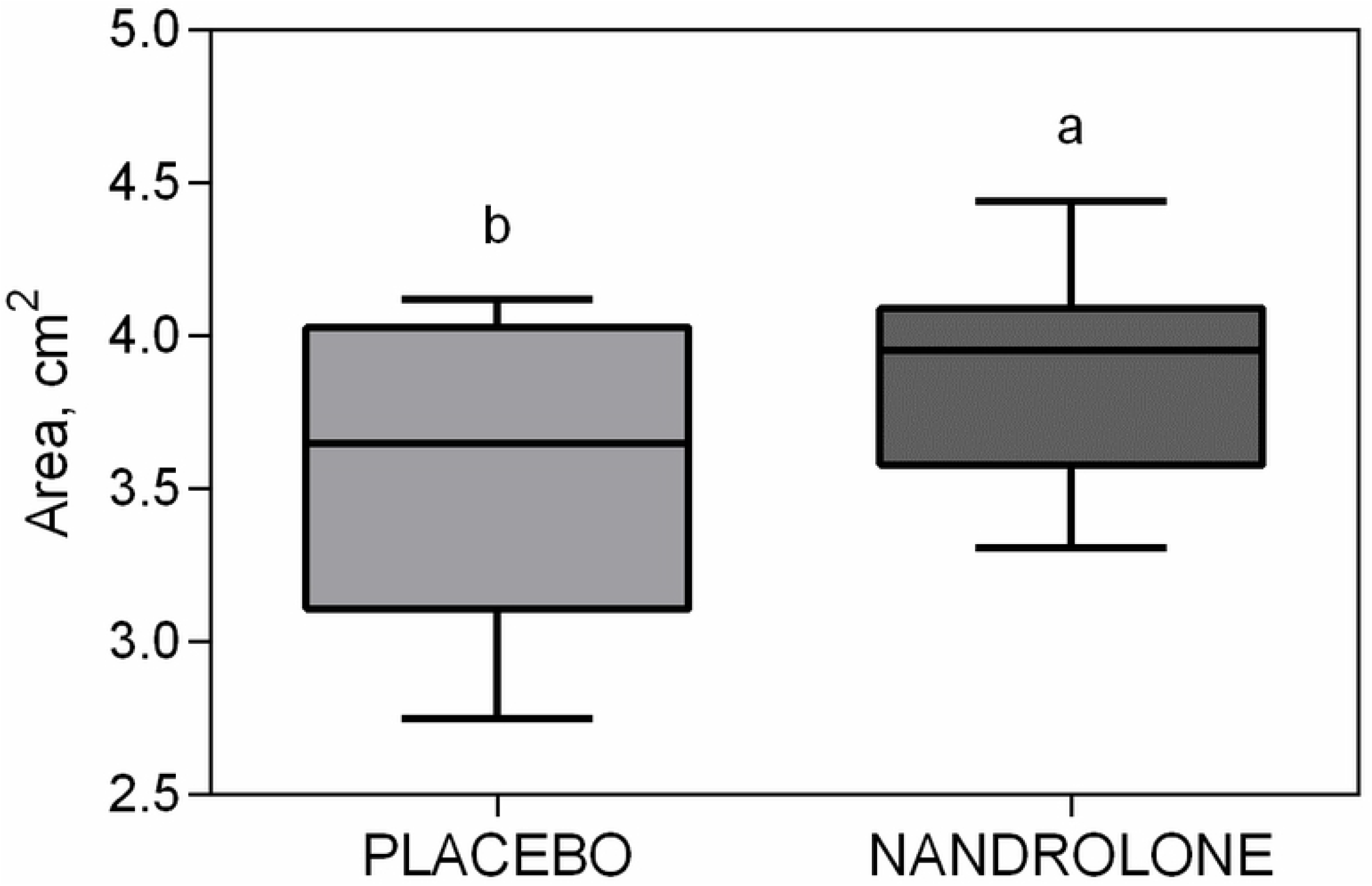
Mean ± SD of Bone area in Placebo and Nandrolone groups. Different letters indicate significant differences by Tukey test (P= 0.0258)

**Fig. 3).**
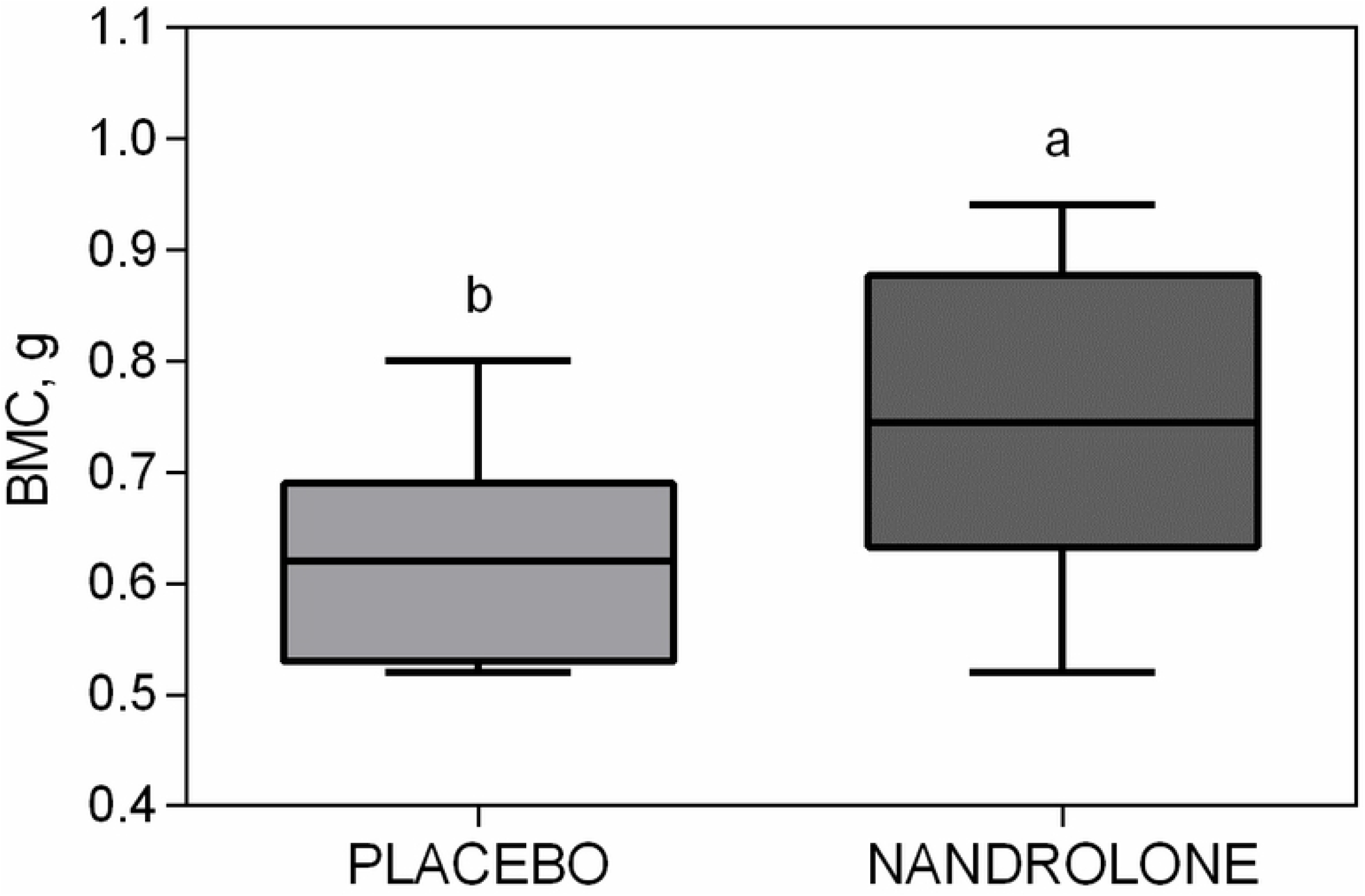
Mean ± SD of BMC in Placebo and Nandrolone groups. Different letters indicate significant differences by Tukey test (P= 0.0140).

Area L was similar between all groups evaluated (P> 0.05). Males had a higher BMC L (P = 0.0317; Fig. 4), and a lower BMD L (P = 0.0468; Fig. 5) than females.

**Fig. 4).**
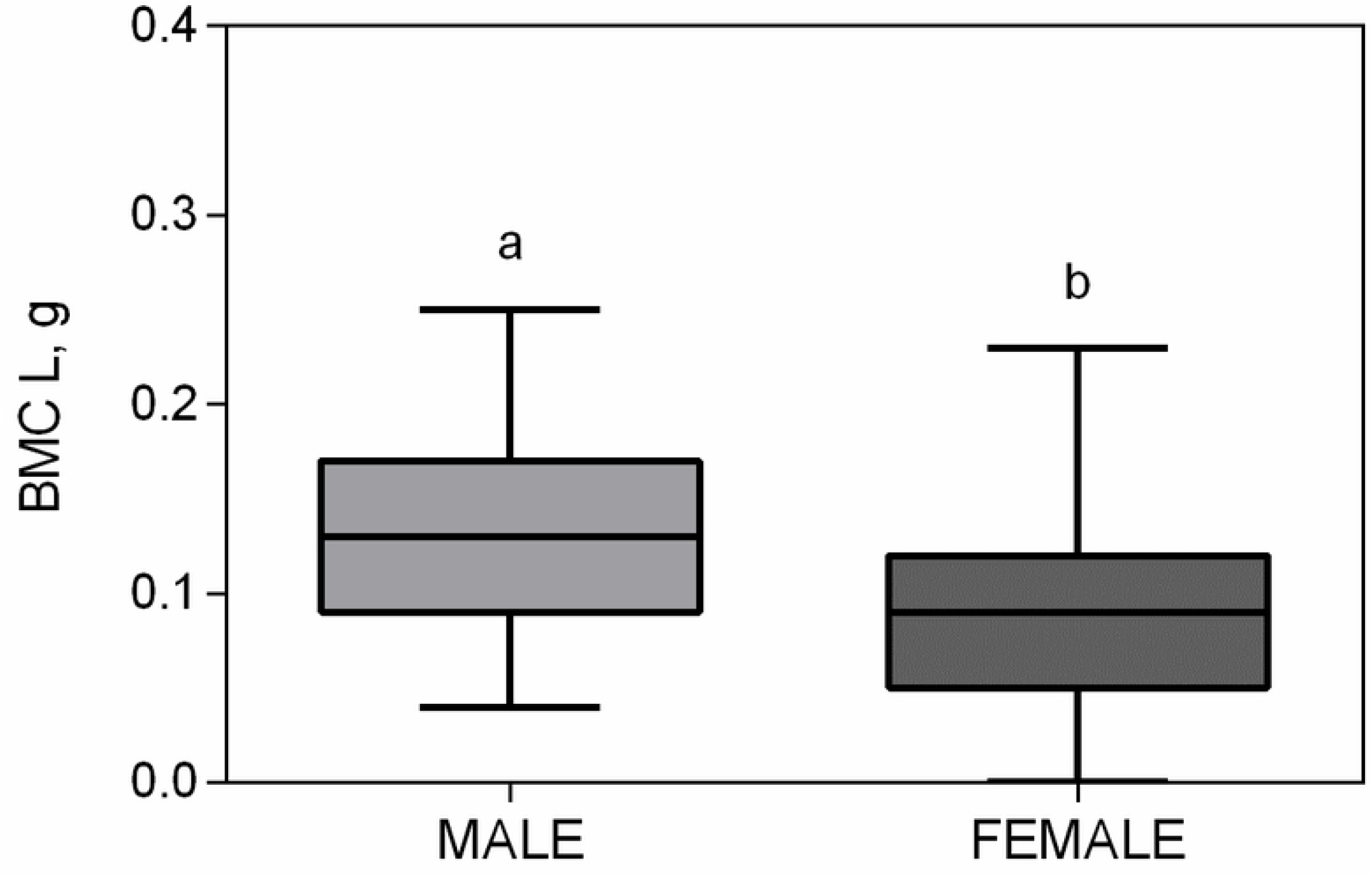
Mean ± SD of BMC L in Male and Female groups. Different letters indicate significant differences by Tukey test (P= 0.0317).

**Fig. 5).**
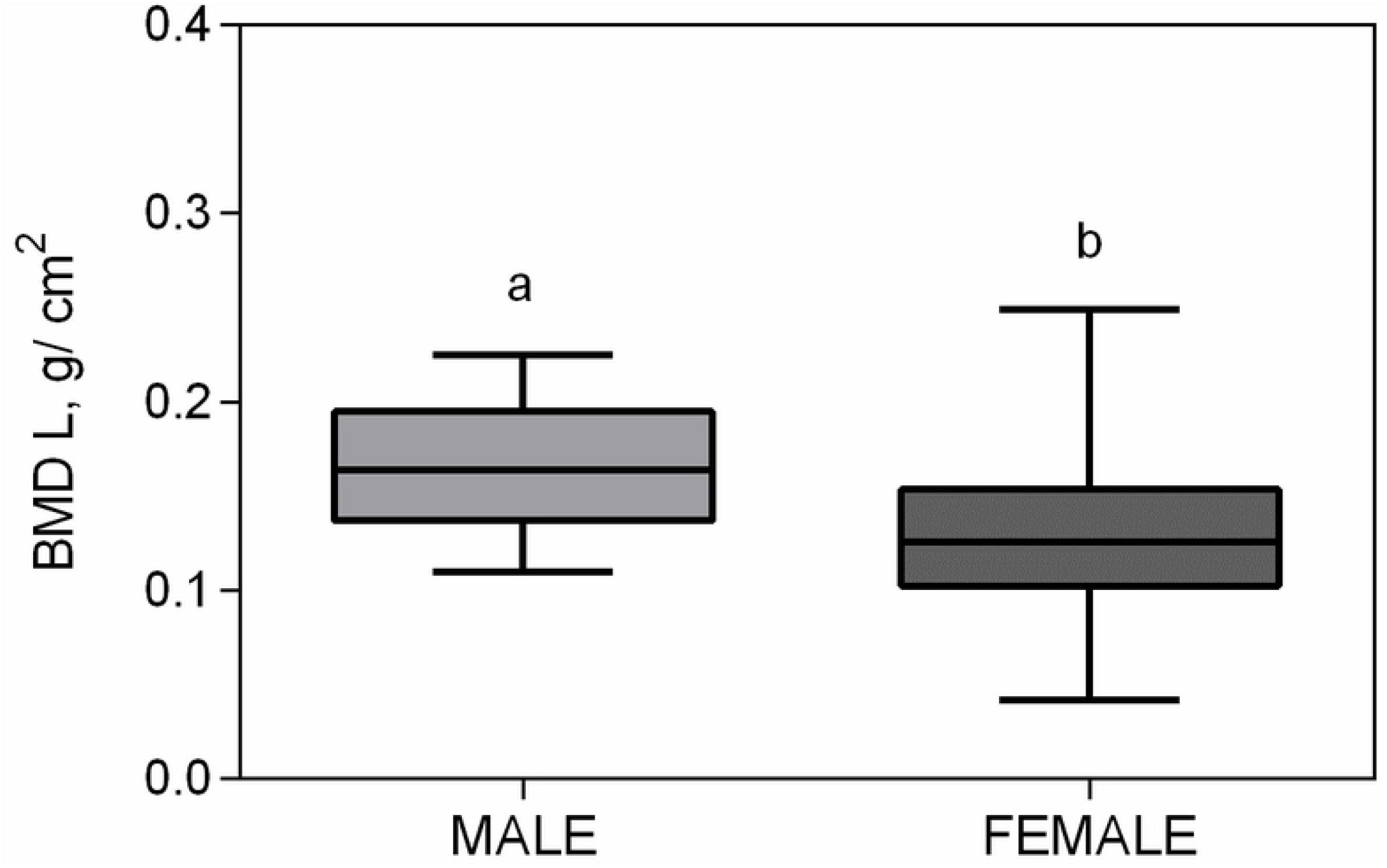
Mean ± SD of BMD L in Male and Female groups. Different letters indicate significant differences by Tukey test (P= 0.0468).

The maturation of the newly formed bone tissue and the presence of medullary spaces was similar between all groups evaluated (Table 3; P> 0.05). However, the Placebo males had higher values on the Extension scale of newly formed bone tissue, when compared to the other groups (P = 0.043). Males treated with Nandrolone showed a higher % of Cartilage presence (5/8) when compared to the other groups (P = 0.006).

**Table 3.** Median ± IQR of bone histologic evaluation in rabbits treated with ND versus placebo.

|  | Placebo |  | Nandrolone |  | P-value |  |  |
| --- | --- | --- | --- | --- | --- | --- | --- |
|  | Male | Female | Male | Female | G | T | G×T |
| Maturation of newly formed bone tissue * | 4.0±0.0 | 4.0±0.0 | 4.0±0.0 | 5.0±1.0 | 0.853 | 0.902 | 0.061 |
| Extension of newly formed bone tissue ** | 3.0±1.0<sup>b</sup> | 5.0±2.5<sup>a</sup> | 5.0±0.0<sup>a</sup> | 5.0±2.5<sup>a</sup> | 0.334 | 0.317 | 0.043 |
| Prevalence of bone tissue *** | 1.0±0.0 | 1.0±0.25 | 1.0±0.0 | 2.0±1.00 | 0.157 | 0.895 | 0.111 |
| Cartilage,% | 0.0 (0/7)<sup>b</sup> | 12.5 (1/7)<sup>b</sup> | 62.5 (5/8)<sup>a</sup> | 0.0 (0/8)<sup>b</sup> | 0.171 | 0.201 | 0.006 |
| Medullary spaces,% | 71.4 (5/7) | 50.0 (4/8) | 37.5 (3/8) | 62.5 (5/8) | 0.285 | 0.855 | 0.569 |

G = male vs female effect evaluated by Friedman test. T = Placebo vs Nandrolone effect evaluated by Friedman test. G × T = Interaction sex and treatment effect evaluated by Kruskal-Wallis test. Different letters in same row indicate significant differences by Dunn's test ( $p < 0.05$ ). \* = evaluated according to the scale where 0 = no periosteal reaction, 1 = moderate periosteal response, 2 = intense periosteal response, 3 = disorganized newly formed bone tissue, 4 = Newly formed bone tissue with moderate organization, and 5 = tissue bone formation with advanced organization. \*\* = assessed according to a scale where 0 = no new bone formation or it was not possible to assess, 1 = mild bone formation, 2 = mild / moderate bone formation, 3 = moderate bone formation, 4 = moderate / intense bone formation, and 5 = intense new bone formation. \*\*\* = assessed according to the scale where 1 = prevalence of primary bone tissue, and 2 = prevalence of secondary bone tissue.

The female placebo group (PG) demonstrated BF with a predominance of primary bone tissue in half of the animals. Only a few animals in the treated group showed a predominance of primary bone tissue in the bone callus region. Treatment with ND in females resulted in better lamellar organization of the secondary bone (Fig. 6). The medullary canal, in the central region of the osteotomy, was similar to a normal medullary canal in a few animals.

**Fig 6:**
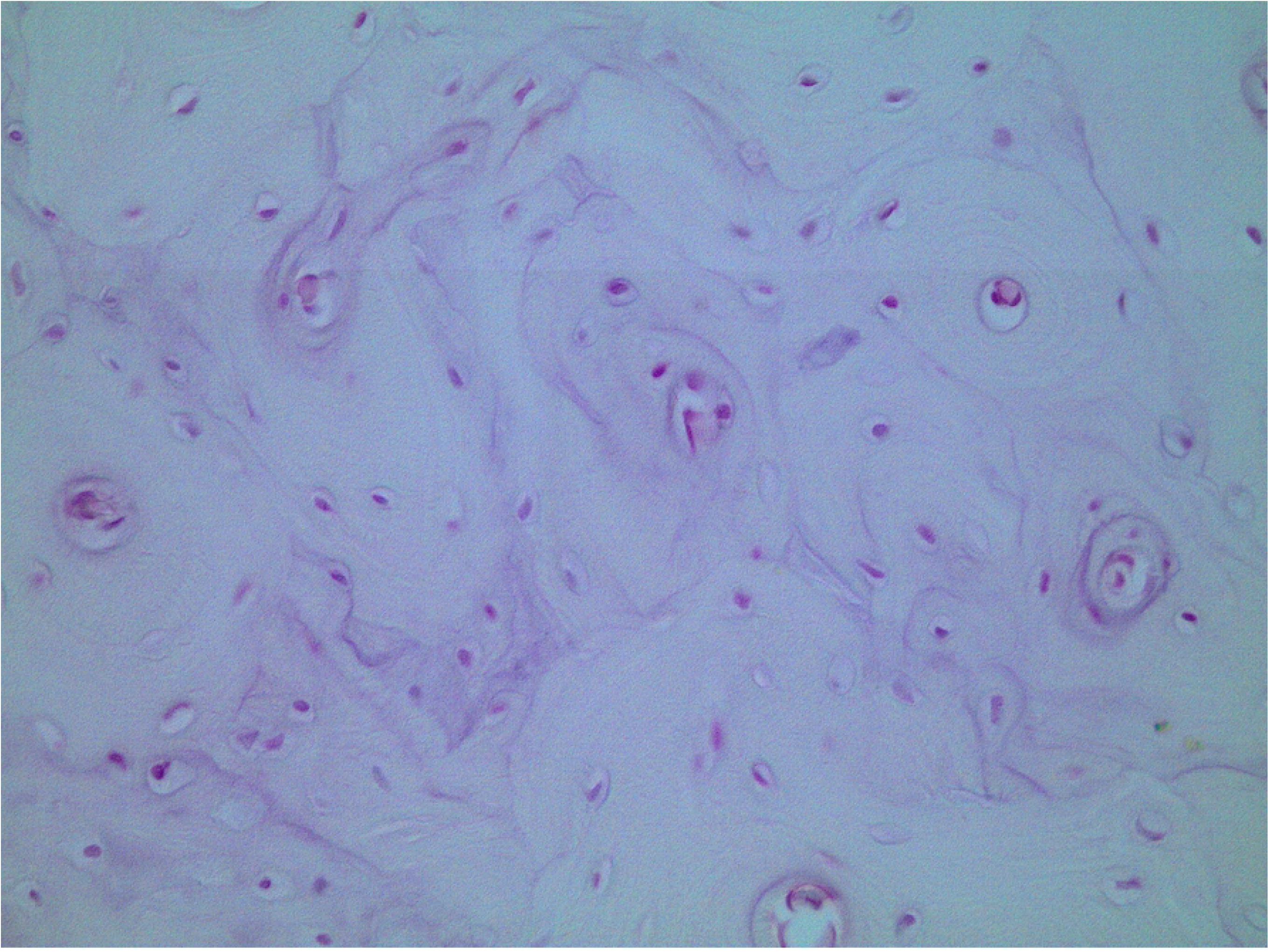
Female Nandrolone 3 - new bone formation at 90 days. There are parallel lamellae arranged on concentric circles around the Haversian canals (mature or secondary bone). HE, 40x lens.

Discrete focal areas of primary bone tissue were observed in most males in the placebo group and focal areas of primary bone tissue, sometimes with fibrocartilage, in the nandrolone group.

## DISCUSSION

There is little published information on the effect of ND on bone healing in experimental and clinical veterinary patients, especially those with nonunion. This study describes some benefits of the use of ND for bone regeneration at the site of critical-size radial defects in rabbits.

The radiographic and densitometric results from this study suggest that ND accelerates the stages of bone consolidation and improves osteoblastogenesis. It also appears to promote an increase in mineral content and bone quality as well as bone filling in critical defects. Previous studies have successfully demonstrated complete filling of bone defects in animals receiving ND, which confirms the acceleration of bone consolidation (5); however, in these studies non-critical circular bone failures of up to 4 mm in diameter were used. It is believed that the experimental failure model used in this study more accurately simulates atrophic bone nonunions. Although complete healing was not achieved in this study the efficacy of the drug in the formation of bone tissue was more effectively evaluated.

In previous studies on ND in guinea pigs, its effectiveness in bone activity was observed; however, these studies are limited because they were evaluated qualitatively. In this study, the evaluations were scored to achieve greater accuracy and reliability of the results (5, 7).

In addition to evaluating the effect of ND on atrophic nonunion, we hypothesised that androgenic therapies might be affected by the physiological and hormonal differences between the sexes. The results obtained in this study suggest that the level of serum circulating testosterone in males may positively affect bone regeneration capacity. There are few reports on this subject in the literature. Previous studies have not demonstrated significant differences between the sexes in young animals (11). Future studies should evaluate the aforementioned parameters and correlate them with the capacity for osteogenesis.

A study by Li et al (12) evaluated the effectiveness of ND against bone loss as an androgenic agent, comparing the use of ND in ovariectomized rats This demonstrated a greater BMD in the evaluated bones in the treated group. In the general densitometric analysis, the female placebo group had lower BMD. This result confirms the effectiveness of ND and suggests that androgens are essential for optimal bone healing.

The same finding was observed in studies that involved osteoporotic women treated with ND. In another study, men showed a 50% increase in the risk of pathological fractures resulting from osteopenia following orchiectomy for gonadal tumours when compared to those who had not undergone the procedure. No decrease in BMD on bone densitometry was seen in men who received androgen therapy in contrast to untreated men (13, 14).

The animals in the nandrolone group showed a greater BMC then the placebo group. Flicker et al. (15) observed a significant increase in bone mineral content in osteopenic females treated with ND, and considering the potential physiological differences of the species, it is suggested that treatment with ND increases bone mineral content, regardless of sex, in rabbits. Guimarães et al. (16) evaluated the influence of ND on bone quality through the evaluation of BMD in male rats after 28 days of treatment and found no significant differences between the treated and control groups. From this, we derive another hypothesis that the beneficial effects of ND may occur later than expected. In our study, we observed any improvement in bone quality from the thirtieth day of treatment.

The microscopic findings showed that the animals in placebo groups had a predominance of primary bone tissue. On the other hand, lamellar organizations of secondary bone and the presence of fibrocartilage were observed in the treated groups. Recently, Senos et al. and Ahmad et al. (17, 7) administered ND for fractures of the femur and mandible in rabbits, and they observed an increase in bone mass, regular bone surface, and reduction in the time to bone regeneration. These results may indicate that ND facilitates osteogenic activity and can accelerate the stages of bone healing.

The degree of fibrocartilaginous tissue formation was different for males and females. This effect of androgens may be important in males, since they are present in higher concentrations and contribute to greater periosteal bone formation and, therefore, greater bone dimensions (13, 18). This is corroborated in this study by the higher percentage of fibrocartilaginous tissue in the NMG. The extent of direct interaction between androgens and osteoblasts or their precursors is still unclear. Androgens affect skeletal growth, including the increase in vascular and muscle cell activity (13).

In this study, treatment with ND resulted in improvement in bone healing in nonunion fractures. However, complete healing did not occur in the NG or PG, as expected, due to the restricted duration of the study. The most important limitation in this study was the inability to assess the side effects of ND, as several studies have shown that the androgen overdose associated with long-term treatment is associated with systemic changes (19–20). Although we did not specifically investigate potential side effects, the animals remained healthy throughout the evaluation. Further studies using ND for a longer period in cases of nonunion of fractures are recommended to assess long-term side effects and the prolonged effect of ND on bone regeneration.

Treatment with ND produced an increase in bone regeneration after the creation of a large defect in the tibia of rabbits. However, this therapy must be combined with surgical treatment, such as fracture fixation and the use of autologous grafts.

## ACKNOWLEDGEMENTS

We thank Professor Dr. Alan Rodrigo Panosso, from the Department of Engineering and Exact Sciences for performing the statistical analyses of this study. We also thank Professor Dra. Silvana Martinez Baraldi Artoni and Professor Dra Lizandra Amoroso from the Department of Animal Morphology and Physiology, who allowed bone densitometry examinations. We are also grateful to CAPES for the funding and support of this research.

## Notes

**Funding:** This study was partly funded by the Coordenação de Aperfeiçoamento de Pessoal de Nível Superior (CAPES) of the Brazilian Government master’s degree scholarship program.

**Conflicts of interest:** The authors report no conflicts of interest.

